# Phase transitions in evolutionary dynamics

**DOI:** 10.1101/361337

**Authors:** Fernando Alcalde Cuesta, Pablo González Sequeiros, Álvaro Lozano Rojo

**Affiliations:** Departamento de Matemáticas, Universidade de Santiago de Compostela, E-15782 Santiago de Compostela, Spain; Departamento de Didácticas Aplicadas, Facultade de Formación do Profesorado, Universidade de Santiago de Compostela, Avda. Ramón Ferreiro s/n, E-27002 Lugo, Spain; Centro Universitario de la Defensa, Academia General Militar, Ctra. Huesca s/n. E-50090 Zaragoza, Spain; IUMA, Universidad de Zaragoza, Pedro Cerbuna 12, E-50009 Zaragoza, Spain; GeoDynApp - ECSING Group, Spain

## Abstract

The evolutionary dynamics of a finite population where resident individuals are replaced by mutant ones depends on its spatial structure. The population adopts the form of an undirected graph where the place occupied by each individual is represented by a node and it is bidirectionally linked to the places that can be occupied by its clonal offspring. There are undirected graph structures that act as amplifiers of selection increasing the probability that the offspring of an advantageous mutant spreads through the graph reaching any node. But there also are undirected graph structures acting as suppressors of selection where, on the contrary, the fixation probability of an advantageous mutant is less than that of the same individual placed in a homogeneous population. Here, we show that some undirected graphs exhibit phase transitions between both evolutionary regimes when the mutant fitness varies. Firstly, as was already observed for small order random graphs, we show that most graphs of order 10 or less are amplifiers of selection or suppressors that become amplifiers from a unique transition phase. Secondly, we give examples of amplifiers of order 7 that become suppressors from some critical value. For graphs of order 6 and 7, we apply computer aided techniques to exactly determine their fixation probabilities and then their evolutionary regimes, as well as the exact critical values for which each graph changes its regime. A similar technique is used to explore a general method to suppress selection in bigger orders, namely from 8 to 15, up to some critical fitness value. The analysis of all graphs from order 8 to order 10 reveals a complex and rich evolutionary dynamics, which have not been examined in detail until now, and poses some new challenges in computing fixation probabilities and times of evolutionary graphs.

## Introduction

In recent times the evolutionary theory on graphs has become a key field to understand biological systems. Although evolutionary dynamics has been classically studied for homogeneous populations, there is now a wide interest in the evolution of populations arranged on graphs after mutant spread. The process transforming nodes occupied by residents into nodes occupied by mutants is described by the *Moran model*. Introduced by Moran [1] as the Markov chain that counts the number of invading mutants in a homogeneous population, it was adapted to subdivided population by Maruyama [2,3] and rediscovered by Lieberman et al. [4] for general graphs. For undirected graphs where links have no orientation, mutants will either become extinct or take over the whole population, reaching one of the two absorbing states, *extinction* or *fixation*. The *fixation probability* is the fundamental quantity in the stochastic evolutionary analysis of a finite population.

If the population is homogeneous, at the beginning, one single node is chosen at random to be occupied by a mutant individual among a population of *N* resident individuals. Afterwards, an individual is randomly chosen for reproduction, with probability proportional to its reproductive advantage (1 for residents and *r* ≥ 1 for mutants), and its clonal offspring replaces another individual chosen at random. In this case, the fixation probability is given by

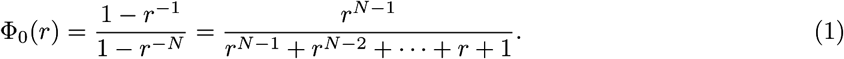

If the population is arranged on nodes of an undirected graph, the replacements are limited to the nodes that are connected by links. According to the Isothermal Theorem [2-4], the fixation probability Φ(*r*) = Φ_0_(*r*) if and only if the graph is *isothermal* (i.e. the temperature *T* = ∑_*j*~*i*_ 1/*d_j_* of any node *i* is constant, where *j* is a neighbor of *i* and *d_j_* is the number of neighbors of *j*), or equivalently *regular* (i.e. the degree *d_i_* of any node *i* is constant). But there are graph structures altering substantially the fixation chances of mutant individuals depending on their fitness. As showed in [4], there are graph structures that *amplify* this advantage. This means the fixation probability function Φ(*r*) > Φ_0_(*r*) for all *r* > 1 for the same order *N*. Notice that Φ(1) = 1/*N* and the inequality must be reversed for *r* < 1. Due to the exact analytical computation of Φ(*r*) given by Monk et al. [5], it is known that star and complete bipartite graphs are *amplifiers of natural selection* whose fixation functions are bounded from above by

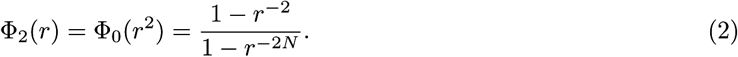

On the other hand, there are also graph structures that *suppress* the reproductive advantage of mutant individuals so that Φ(*r*) < Φ_0_(*r*) for all *r* > 1. Examples of this kind of graph structures were known for some fitness values (see [6]). In [7], we introduced examples of *suppressors of natural selection* of order 6, 8 and 10 (see Fig 1), denoted by *ℓ*_6_, *ℓ*_8_ and *ℓ*_10_, whose fixation probabilities remain smaller than Φ_0_(*r*) for every *r* > 1.

**Fig 1.**
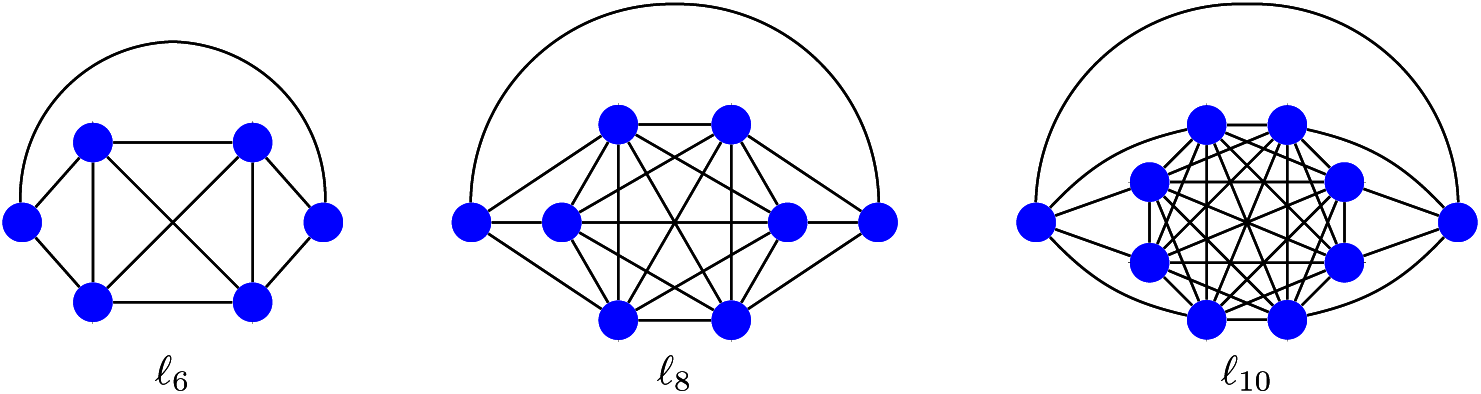
Suppressors of order 6, 8 and 10 for any fitness value. In [7], we called *ℓ*-*graph* the undirected graph of even order *N* = 2*n* + 2 ≥ 6 obtained from the clique *K*_2*n*_ by dividing its vertex set into two halves with *n* ≥ 2 vertices and adding 2 extra vertices. Each of them is connected to one of the halves of *K*_2*n*_ and with the other extra vertex.

The analysis of disadvantageous mutants (with *r* < 1) is also interesting when comparing amplification and suppression of selection for graphs, but here, for simplicity, we focus on the case of advantageous mutant (with *r* > 1). Different initialization and updating types have been also considered in [8] and [9], see also [10] for a comparative analysis of update mechanisms. If the initial distribution is uniform (i.e. the probability that a node will be occupied by the initial mutant is equal for all the nodes) and the graph evolves under Birth-Death updating, Hindersin and Traulsen showed in [9] that almost all small undirected graphs are amplifiers of selection. Assuming both conditions and focusing on the advantageous case, we distinguish two different evolutionary regimes out of the isothermal one: given values 1 ≤ *r*_0_ < *r*_1_ ≤ +∞, a graph is an *amplifier of selection for r* ∅ (*r*_0_, *r*_1_) if the fixation probability function Φ(*r*) > Φ_0_(*r*) and a *suppressor of selection for r* ∅ (*r*_0_, *r*_1_) if Φ(*r*) < Φ_0_(*r*) for all *r*_0_ < *r* < *r*_1_. Star and bipartite complete graphs are amplifiers for *r* ∈ (1, +∞) and graphs *ℓ*_6_, *ℓ*_8_ and *ℓ*_10_ are suppressors for *r* ∈ (1, +∞). There are suppressors that become amplifiers from some critical value *r_c_* > 1 (see Fig 2). Borrowing the terminology from percolation theory, in this case, we say *r_c_* is a *transition phase* from the suppressor regime to the amplifier regime. In general, we say that a number *r_c_* > 1 is *a transition phase* between two different evolutionary regimes if there are values 1 ≤ *r*_0_ < *r*_1_ ≤ +∞ such that the graph is a suppressor (resp. an amplifier) for *r* ∈ (*r*_0_, *r_c_*) and an amplifier (resp. a suppressor) for *r* ∈ (*r_c_*,*r*_1_).

**Fig 2.**
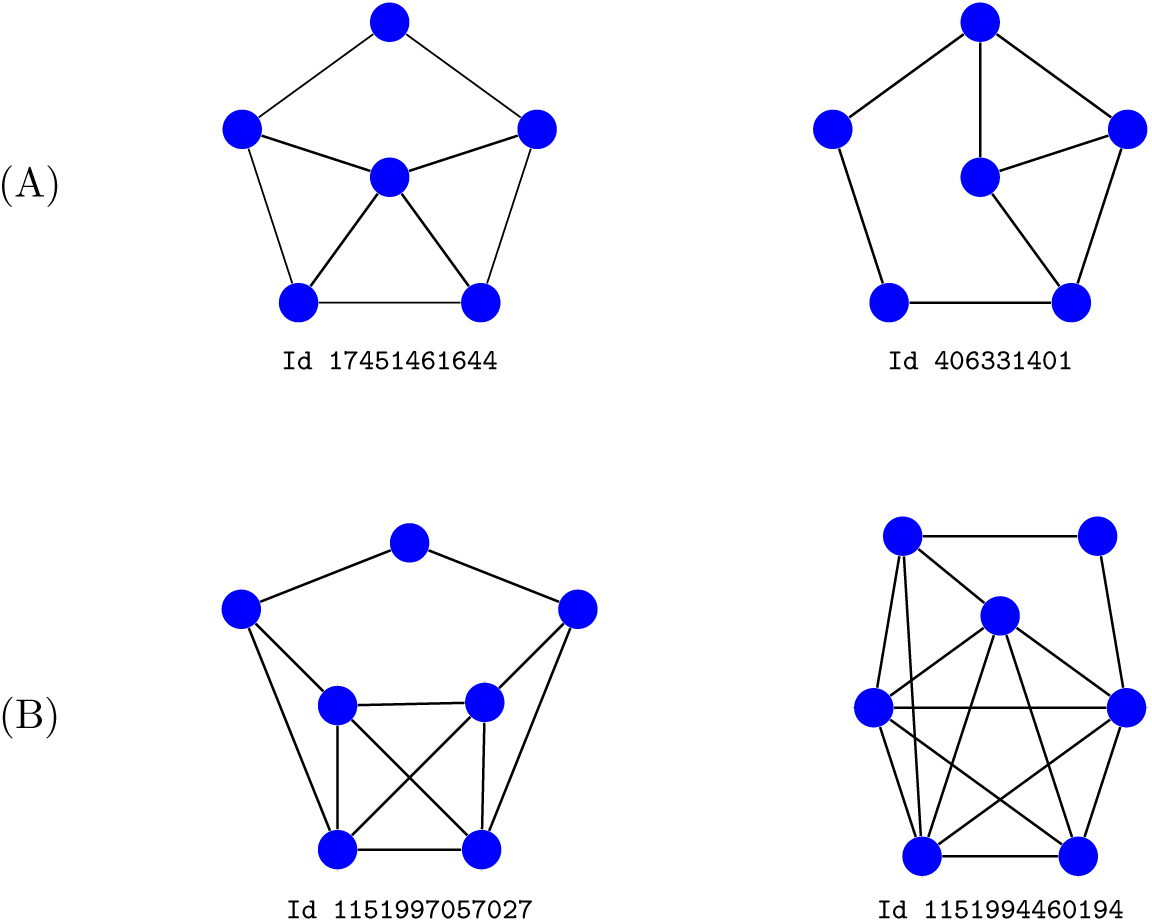
Suppressors that become amplifiers. (A) Examples of order 6 from [11] with a unique transition phase at *r_c_* ≈ 2.82 and *r_c_* ≈ 4.56 respectively. (B) Examples of order 7 from [6] having a unique transition phase at *r_c_* ≈ 79.15 and *r_c_* ≈ 1.98 respectively. In both cases, we use identification numbers from [11].

In this paper, we show that most graphs of order 10 or less are amplifiers or suppressors that become amplifiers from a unique transition phase *r_c_* > 1. However, we also exhibit two other types of transitions which are much more interesting:

- There are amplifiers of order greater or equal to 7 that become suppressors from a unique transition phase *r_c_* > 1.
- There are graphs of order greater or equal to 8 exhibiting more than one transition phase.

Since the fixation probabilities Φ_0_(*r*) and Φ(*r*) are monotonically increasing, these results were not exactly expected, while numerical simulations should be considered with caution. In fact, for finite populations, it has been suggested that results obtained for weak selection may remain valid when the selection is no longer weak. However, in [12], Wu et al. showed that this is no the case for homogeneous populations under frequency dependent selection. Here, we show that the graph structure can completely alter the evolutionary regime of a structured population even under constant selection. In addition, the fact that the survival chances of mutant individuals may decrease with respect to a homogeneous population when their fitness increases might have interesting biological consequences. Our results also demonstrate that numerical simulation is not enough to determine the evolutionary regime of a graph for the whole range of fitness values. Moreover, the very slight difference between different regimes for some graphs often forces to consider an extremely high number of trials when using Monte Carlo method, like recently in [13] to show a family of graphs of order ≥ 12 that changes from amplifier to suppressor as *r* increases. For example, 10^10^ trials were needed in [7] to determine the evolutionary regime of the graphs *ℓ_N_* for 12 ≤ *N* ≤ 24 and 1 ≤ *r* ≤ 4.

In [11], we presented an extremely precise database of fixation probabilities for all undirected graphs of order 10 or less for fitness values *r* varying from 0.25 to 10 with step size of 0.25 (see details in Materials and Methods). Here, we use this database to detect new evolutionary dynamics: in this way, we identify two amplifiers of order 7 with a transition to the suppressor regime at some *r_c_* ≤ 10. Then we adapt the technique described in [7] to symbolically compute the exact fixation probability Φ(*r*) of these examples for all *r* > 1 proving that they really have a unique transition phase. A similar procedure is used to exactly determine the evolutionary regime of any graph of order 7 or less for any fitness value. Thus, we find a third example of order 7 exhibiting a transition from the amplifier to the suppressor regime for *r_c_* > 10. The first method is also applied to show the existence of graphs of order 8, 9 and 10 with more than one transition phase. This reveals that undirected graphs have a complex and rich evolutionary dynamics that are worth studying in detail.

## Materials and Methods

### Mathematical model

Let *G* be an undirected graph with node set *V* = {1, …, *N*}. In fact, all graphs considered here will be assumed connected. Denote by *d_i_* the degree of the node *i*. The *Moran process* on *G* is a Markov chain *X_n_* whose states are the sets of nodes *S* inhabited by mutant individuals at each time step *n*. The transition probabilities are obtained from the matrix *W* = (*w_ij_*) given by *W_ij_* = 1/*d_i_* if *i* and *j* are neighbors and *W_ij_* = 0 otherwise. More precisely, for each fitness value *r* > 0, the transition probability between *S* and *S′* is given by

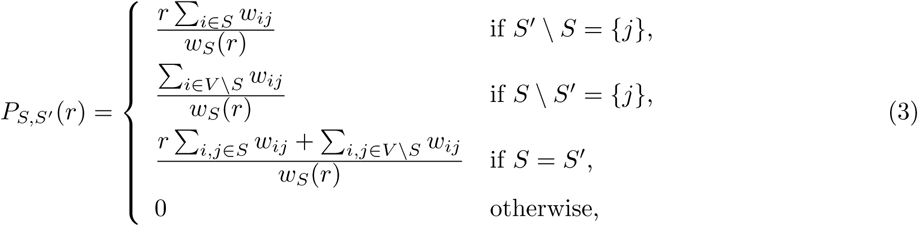

where

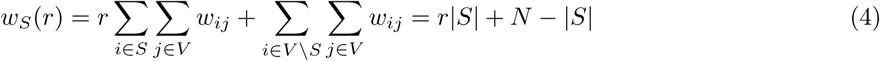

is the total reproductive weight of resident and mutant individuals. The *fixation probability* of each subset *S* ⊂ *V* inhabited by mutant individuals

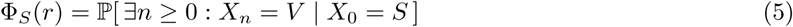

is a solution of the system of 2^*N*^ linear equations

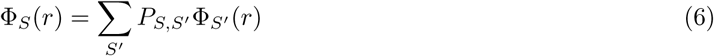

with boundary conditions Φ_∅_(*r*) = 0 and Φ_*v*_ (*r*) = 1. The *(average) fixation probability* is given by

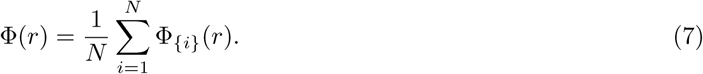

Denoting by ***P***(*r*) = (*P*_*s*,*s′*_(*r*)) the transition matrix, Eq 6 can be written as

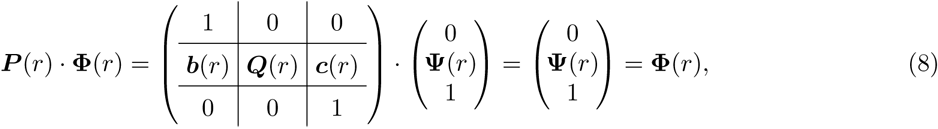

where **Φ**(*r*) = (0, **Ψ**(*r*), 1) is the vector with coordinates Φ_*S*_(*r*), (1, ***b***(*r*), 0) is the vector with coordinates *P*_*S*,∅_(*r*), and (0, ***c***(*r*), 1) is the vector with coordinates *P*_*S*,*V*_(*r*). It can be also rewritten as

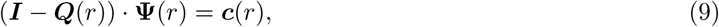

where ***I*** is the identity matrix of size 2^*N*^ − 2. This equation has a unique solution **Ψ**(*r*) whose coordinates are rational functions on *r* with rational coefficients [7, S1 Text]. But considering Eq 3, we can multiply the equation associated to each state *S* by *w_S_* (*r*) reducing Eq 9 to

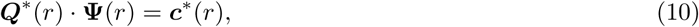

where the entries of ***Q***^∗^(*r*) and ***c***^∗^ (*r*) are now degree one polynomials with rational coefficients.

### Database

In [11], we presented an accurate database of the fixation probabilities for all connected undirected graphs with 10 or less vertices, which means 11,989,763 graphs excluding the trivial one with one single vertex. The generation of the edge lists was done with SageMath, whereas the computation of Φ(*r*) was written in the C programming language. Firstly, we compute the matrix ***Q***^∗^ (*r*) and the vector ***c***^∗^ (*r*) whose entries are represented as couples *a* and *b* of pairs of 64 bits integers. Next, we solve Eq 10 for each fitness value *r* varying from 0.25 to 10 with step size of 0.25. Each graph is identified (up to isomorphism) with a unique 64 bits unsigned integer, which allows us to recover the edge list without previous knowledge of its order or size, see again [11] and references therein for details. The database is presented as a single HDF5 file available at [14].

### Computation method

A method to compute the exact (average) fixation probability Φ(*r*) of graphs of small order with symmetries was described in [7]. As we already said, Φ(*r*) = Φ′(*r*)/Φ″(*r*) is a rational function where the numerator
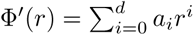
and the denominator
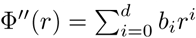
are polynomials with rational coefficients of degree *d* ≤ 2^*N*^ − 2. Symmetries are used to reduce the degree *d* and therefore the number 2(*d* + 1) of coefficients involved in the computation of Φ(*r*). Since Φ(*r*) converges to 1 as *r* → +∞, we have *a_d_* = *b_d_* = 1 and then Eq 6 can be replace with a system of 2*d* linear equations

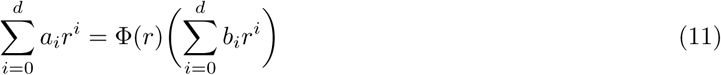

that arise from evaluating Φ(*r*) for fitness values *r* ∈ {1,…, *d* + 1,1/2,…, 1/*d*}. Instead of the SageMath program given in [7], a new C++ program available from S1 File allows us now to symbolically

- compute the fixation probability Φ(*r*) for these values, and
- solve the reduced linear system Eq 11.

Once Φ(*r*) has been calculated solving this system, we can determine the states and the phase transitions of the graph by computing the sign and the zeros of the rational function Φ(*r*) – Φ_0_(*r*).

## Results

Contrary to a widespread idea that it is easier to construct suppressors of selection than amplifiers of selection, which is true only if the edges are oriented, Hindersin and Traulsen [9] showed that most undirected (Erdös-Rényi) random graphs of small order are amplifiers of selection under Birth-Death updating. Here, by regarding all graphs of order 10 or less, we confirm that most undirected graphs of order ≤ 10 are amplifiers of selection for fitness values *r* ≤ 10, see Table 1. In order 6, there are exactly one suppressor of selection, namely the graph *ℓ*_6_ described in [7] (see Fig 1), five isothermal graphs, six suppressors that become amplifiers from a unique critical value, and the remaining 100 are amplifiers of selection (see S1 Table). Two of suppressors changing into amplifiers was already described in [11] (see Fig 2.(A)). All the graphs portrayed in the paper are gathered in S1 Fig and S2 Fig with indication of their identification numbers, names, regimes and transitions.

**Table 1.** Number and percentage of isothermal and suppressor graphs, as well as graphs exhibiting one or more phase transitions. Graphs are determined up to isomorphism, so any graph cannot be mapped to each other via a permutation of vertices and edges. For graphs of order 8 and more, we distinguished between suppressors and apparent suppressors as fitness values vary only between 1 and 10. However, additional exact computations for some higher values of the fitness *r* (varying from 10 to 2, 000 with step size of 1) seem exclude more than two transitions in order 8 and 9 and more than three transitions in order 10. Exact results are marked in bold: a complete description of the phase transitions for graphs of order 6 and 7 is given in S1 Table.

| N | # | Iso | Sup | 1 Trans | 2 Trans | 3 Trans |
| --- | --- | --- | --- | --- | --- | --- |
| 6 | 112 | 5 (4.46%) | 1 (0.89%) | 6 S/A (5.36%) |  |  |
| 7 | 853 | 4 (0.47%) | 3 (0.35%) | 52 S/A (6.10%)<br>3 A/S (0.35%) |  |  |
| 8 | 11,117 | 17 (0.15%) | 90 (0.81%) | 427 S/A (3.84%)<br>36 A/S (0.32%) | 3 S/A/S (0.03%) |  |
| 9 | 261,080 | 11 (<0.01%) | 1,951 (0.75%) | 9,489 S/A (3.63%)<br>854 A/S (0.33%) | 43 S/A/S (0.02%)<br>6 A/S/A (<0.01%) |  |
| 10 | 11,716,571 | 167 (<0.01%) | 91,110 (0.78%) | 407,001 S/A (3.47%)<br>40,974 A/S (0.35%) | 3,086 S/A/S (0.03%)<br>578 A/S/A (<0.01%) | 19 S/A/S/A (<0.01%) |

N = Order, # = Number, **Isotherm** = Isothermal, **Sup** = Suppressor, **Trans** = Transition, **S/A** = Transition from suppressor to amplifier, **A/S** = Transition from amplifier to suppressor, **S/A/S** = Double transition Suppressor/Amplifier/Suppressor, **A/S/A** = Double transition Amplifier/Suppressor/Amplifier, **S/A/S/A** = Triple transition Suppressor/Amplifier/Suppressor/Amplifier.

### Amplifiers and suppressors of order 7

A close look to the barcode diagram for the 853 graphs of order 7 shown in Fig 3 reveals a new phenomenon: we distinguish two amplifiers that become suppressors from a critical value *r_c_* ≤ 10. To determine their evolutionary regime and ensure that there are not other transitions, we compute the average fixation probability Φ(*r*) according to the procedure explained in Materials and Methods: we find three suppressors of selection, namely Id 1134281908237, Id 1134281902105, and Id 1151998128135, and a number of suppressors that later become amplifiers of selection, namely 52, including the suppressors presented in [6] and depicted in Fig 2.(B). Moreover, we find indeed three amplifiers for weak selection Id 1151592835082, Id 1151860745228, and Id 1151592837126 with transitions at *r_c_* ≈ 4.98, *r_c_* ≈ 6.37 and *r_c_* ≈ 24.79 respectively. A quantitative resume is given in Table 1, while some graphs are depicted in S1 Fig and S2 Fig.

**Fig 3.**
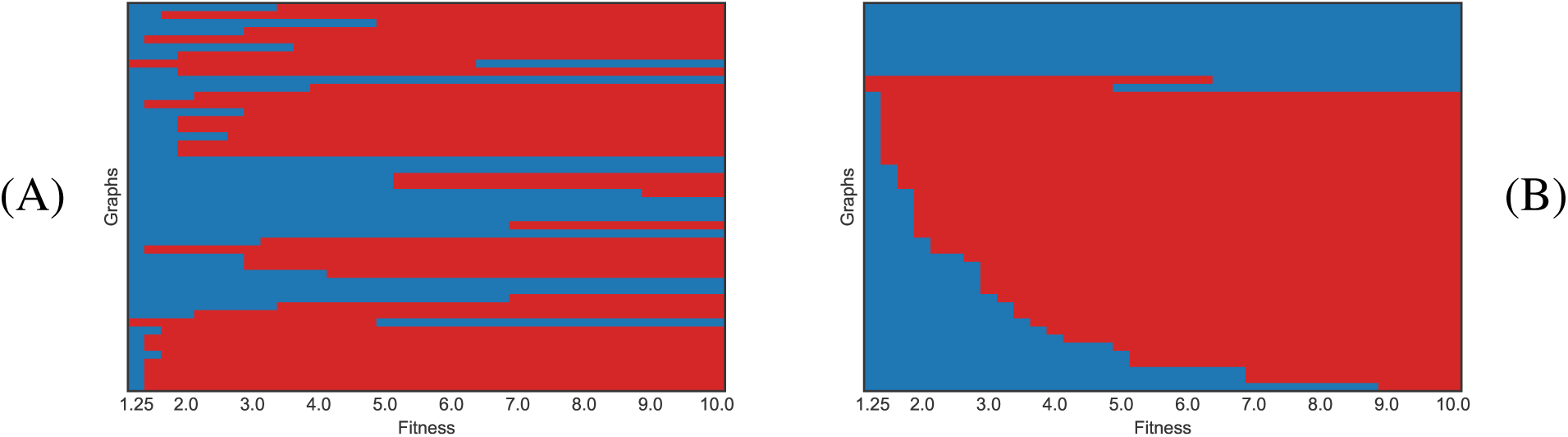
Barcodes describing transition phases of graphs of order 7. Each horizontal line corresponds to a graph, and color represents the evolutionary regime for the given fitness: blue color corresponds to suppressor regime and red color to amplifier regime. (A) Unsorted data for suppressors and graphs with one transition. (B) Sorted data for suppressors and graphs with one transition.

The two first suppressors Id 1134281908237 and Id 1134281902105 are shown in Fig 4, where we also added a third graph Id 1151998648333 with a unique transition from the suppressor to the amplifier regime at *r_c_* ≈ 5.17. Their construction is very similar to *ℓ*-*graphs* defined in [7]. Recall that
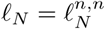
is an undirected graph of even order *N* = 2*n* + 2 ≥ 6 obtained from the clique *K*_2*n*_ by dividing its vertex set into two halves with *n* ≥ 2 vertices and adding 2 extra vertices. Each of them is connected to one of the halves of *K*_2*n*_ and with the other extra vertex (see Fig 1). More generally, we denote by 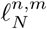
the undirected graph obtained adding two interconnected extra vertices to the clique *K*_*N*−2_ and connecting each one to disjoint families of vertices in the clique having *n* and *m* elements with *n* + *m* ≤ *N*. We say that
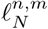
is *balanced* if *n* = *m* and *unbalanced* otherwise. As explained in S1 Fig, using this notation, the third graph in Fig 4 can be simply denoted by
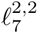.

**Fig 4.**
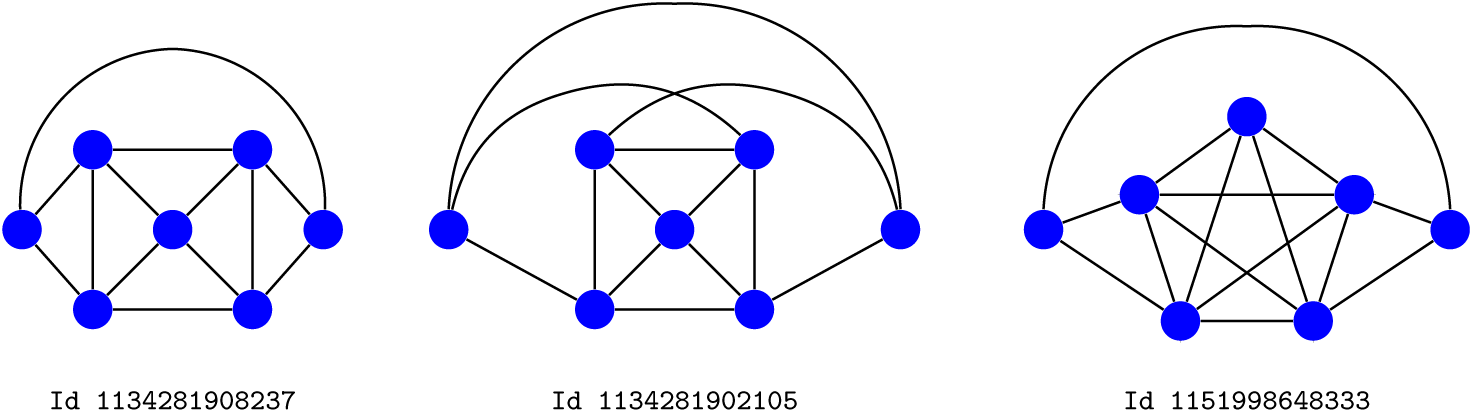
suppressors for weak selection and beyond. Using identification numbers from [11], the graphs Id 1134281908237 and Id 1134281902105 are suppressors for any fitness value, while Id 1151998648333 exhibit a unique transition suppressor/Amplifier at *r_c_* ≈ 5.17.

### Phase transitions for *ℓ*-graphs

For the three graphs in Fig 4, the suppression mechanism seems directly related to the suppressor nature of *ℓ*-graphs [7] and *clique*-*wheels* [15]. Indeed, on a star graph, it is more likely that a peripheral mutant survives and reproduces occupying the central node. On the contrary, on a complete graph, it is much more unlikely that the initial mutant will survive surrounded by residents. But in *ℓ*-graphs and clique-wheels, the survival chances of the mutants placed at the central clique are reduced by its connection with peripheral nodes occupied by residents, as well those of the peripheral mutants connected with central residents, although a subtle balance seems to be needed to suppress the reproductive advantage of mutant individuals according to our data. To confirm this idea, we explore evolutionary regimes and transitions of some balanced and unbalanced *ℓ*-graphs.

Unbalanced *ℓ*-graphs, namely
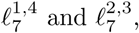
were studied in [7] using Monte Carlo simulation. Both were shown as suppressors; the first one was shown as amplifier from a relatively small fitness value, whereas the second one was shown as suppressor for any fitness value *r* ≤ 10. Now we know they exhibit a unique transition phase (from the suppressor to the amplifier regime) at *r_c_* ≈ 1.80 and *r_c_* ≈ 25.47 respectively.

In order 8, we consider the graphs
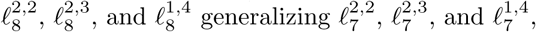
which are depicted in Fig 5. They still are suppressors that become amplifiers from critical values *r_c_* ≈ 4.15, *r_c_* ≈ 5.32 and *r_c_* ≈ 1.89 respectively.

**Fig 5.**
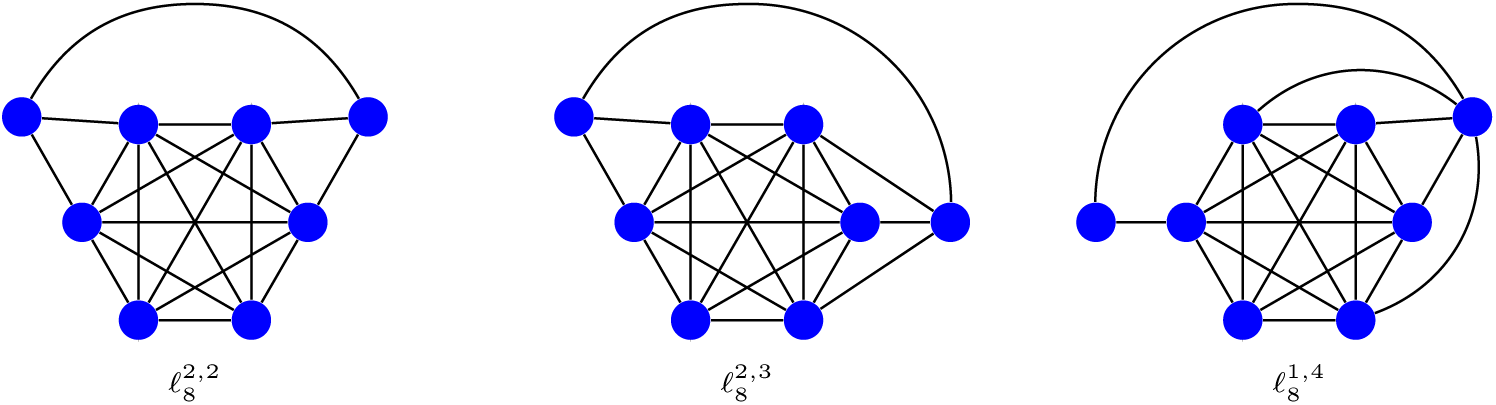
*ℓ*-graphs of order 8. The graphs
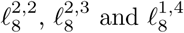
(with identification numbers Id 38605195624473, Id 38605195624473 and Id 38605187250242) have a unique transition of type suppressor/Amplifier at *r_c_* ≈ 4.15, *r_c_* ≈ 5.32 and *r_c_* ≈ 1.89 respectively.

Due to the symmetries of the graphs
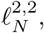
our computation method is also applicable to these graphs when *N* varies from 6 to 15. The exact differences Φ_0_(*r*) – Φ(*r*) are depicted in Fig 6 when *r* varies from 1 to 10, although the monotonous behavior of transitions can be better seen in S2 Table. As before, all these graphs are suppressor with a unique transition phase to the amplifier regime.

**Fig 6.**
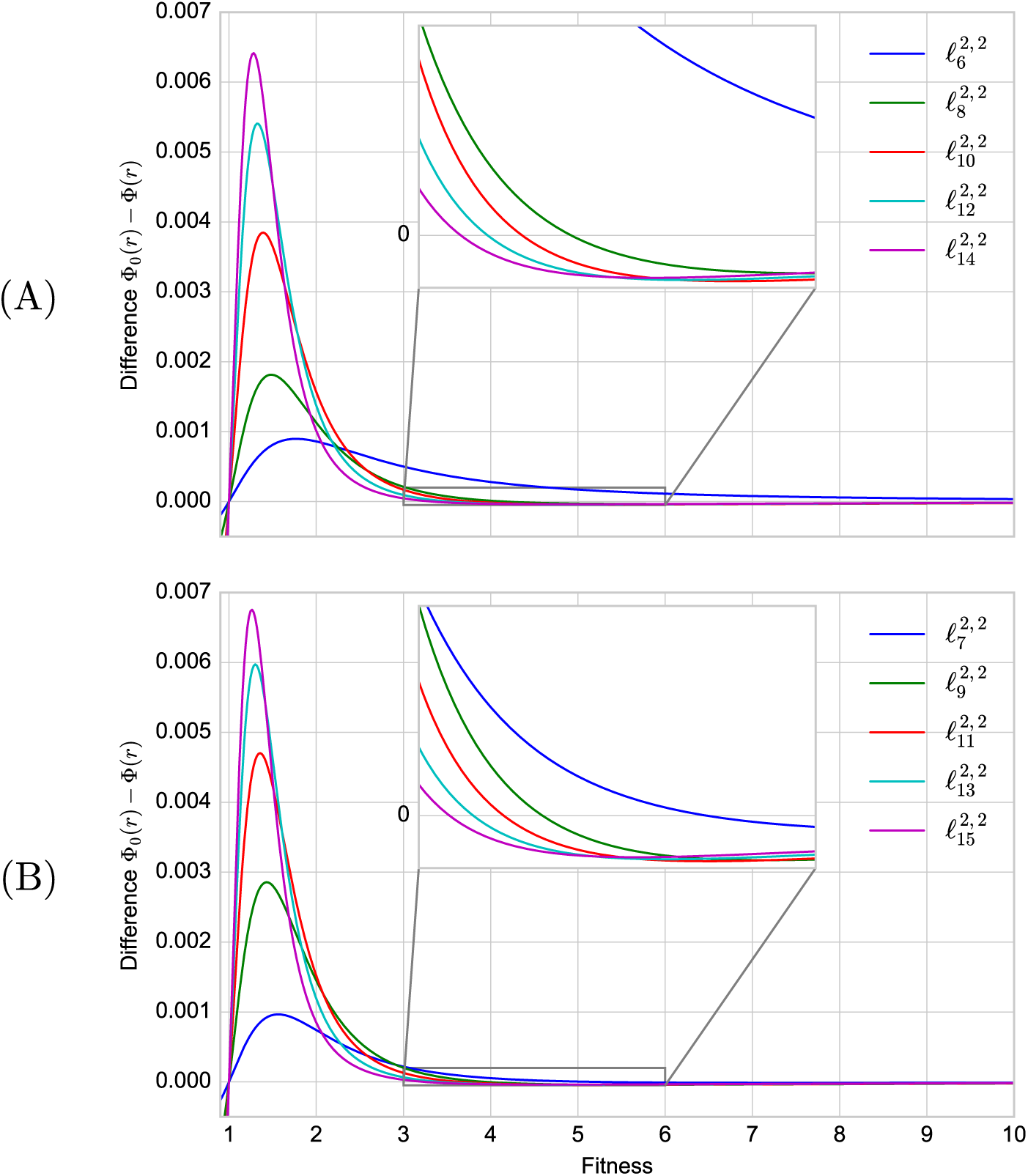
The exact differences Φ_0_(*r*) − Φ(*r*) for the graphs
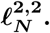. (A) Even orders. (B) Odd orders. Initial evolutionary regimes are distinguishable from the graphic of functions Φ_0_(*r*) − Φ(*r*) associated to the graphs
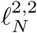
when *N* varies from 6 to 15, while transitions can be observed in the zoomed images. The exact phase transitions are specified in S2 Table.

### Transitions from amplifier to suppressor regime

As we have already said, there are three amplifiers Id 1151592835082, Id 1151860745228, and Id 1151592837126 that become suppressors from critical values *r_c_* ≈ 4.98, *r_c_* ≈ 6.37 and *r_c_* ≈ 24.79. The first and the last of these graphs are constructed from the same building blocks (see Fig 7). Thus, we say that Id 1151592835082 is a *friendship cycle*
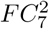
and Id 1151592837126 is a *friendship star*
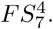
A friendship cycle
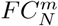
is a cycle *C_n_* where *m* disjoint pairs of neighbors are connected to an extra central vertex so that two vertices have at most one neighbor in common, the total order *N* being equal to *n* +1. In a friendship star
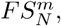
two neighbors can share two different neighbors, but they cannot be connected either to each other, nor to another new neighbor. Additionally, one single vertex can belong to more than two 3-cycles composing a friendship subgraph (to distinguish it from a friendship ribbon
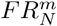
where no vertex can belong to more than two 3-cliques, see S2 Fig). Here *N* is the number of vertices and m is the number of 3-cliques.

**Fig 7.**
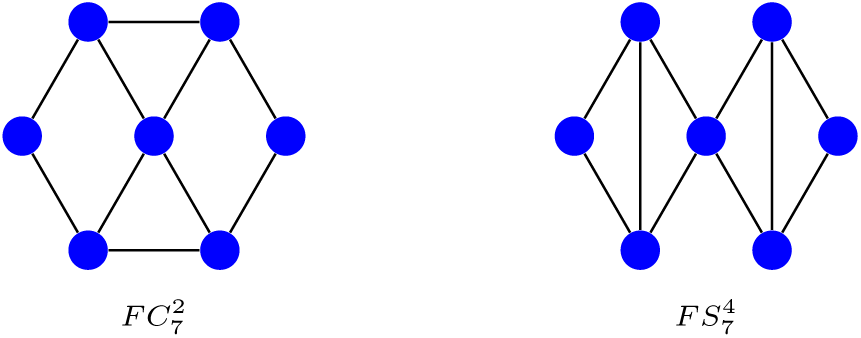
Amplifiers that become suppressors. The friendship cycle
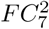
(with identification number Id 1151592835082), which becomes a suppressor from *r_c_* ≈ 4.98, and the friendship star
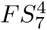
(with identificatin number Id 1151592837126), which becomes a suppressor from *r_c_* ≈ 24.79.

We symbolically compute the fixation probability Φ(*r*) for the friendship cycles
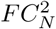
with *N* = 7, 8, 9 proving that they are amplifiers transformed into suppressors from fitness values *r_c_* ≈ 4.98, *r_c_* ≈ 3.12, and *r_c_* ≈ 2.45 respectively (see Fig 8-(A)). For bigger orders *N* varying from 10 to 15, we compute the fixation probabilities for selected fitness values *r* between 1 and 10. The differences Φ_0_(*r*) − Φ(*r*) are shown in Fig 8-(B). Only
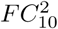
shows a transition in that interval and it is not clear that the others friendship cycles
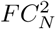
evolve from the amplifier to the suppressor regime. However, the friendship cycle
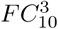
(see S2 Fig) also is an amplifier that becomes a suppressor from *r_c_* ≈ 4.09.

**Fig 8.**
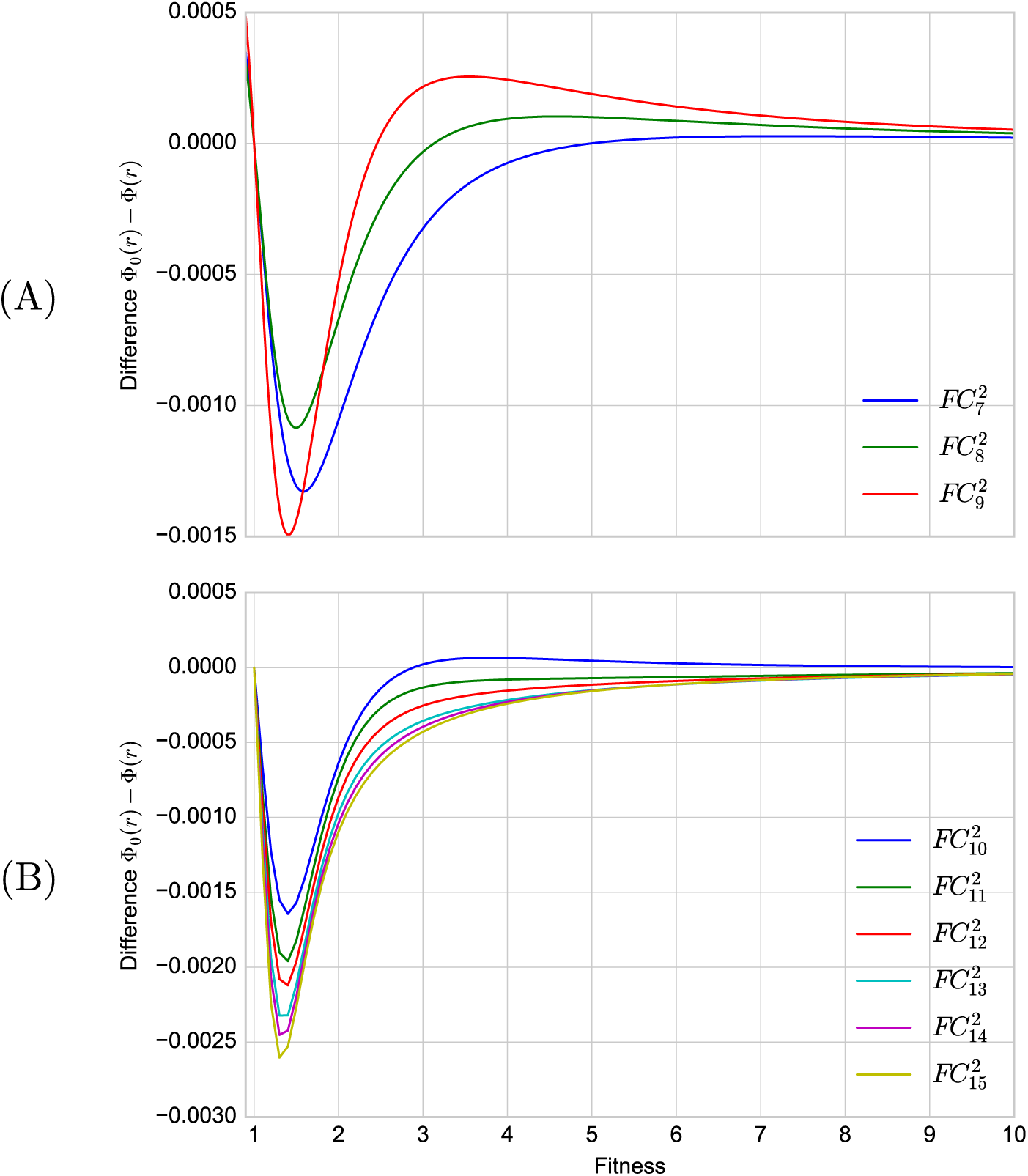
The differences Φ_0_(*r*) − Φ(*r*) for the friendship cycles
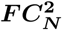
with *N* varying from 7 to 15. (A) The exact differences for *N* =7, 8, 9. (B) The differences computed for 10 ≤ *N* ≤ 15 and 1 ≤ *r* ≤ 10 by combining the method described in [11] with the use of symmetries.

On the other hand, we also symbolically computed the Φ(*r*) for the friendship star
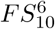
showing that it becomes a suppressor at *r_c_* ≈ 9.96. Since there are not enough symmetries to reduce the system of equations, this approach could not be pushed any further. The same happens with
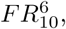
and therefore we have to use the method from [11] seeing that this is an amplifier for *r* ≤ 10. Since we extended the interval until *r* = 2000 without finding any transition, this is probably a global amplifier. These graphs are also depicted in S2 Fig.

### Amplifiers and suppressors of order 8 and more

Other interesting phenomena are also revealed by the barcode diagrams in order 8 and higher as shown in S3 Fig, S4 Fig, and S5 Fig. Now, the number of amplifiers and suppressors is only approximate since the fitness is limited to the interval [1,10]. Thus, we distinguished between suppressors and apparent suppressors, that is, those being suppressors for 1 < *r* ≤ 10. The existence of double transitions of type suppressor/Amplifier/suppressor is clearly visible in S3 Fig. The three graphs of order 8 with a double transition are portrayed in S6 Fig. Though we must add a few transitions of type Amplifier/suppressor/Amplifier for the graphs of order 9, as shown in S4 Fig. A number of double transitions is also shown in S5 Fig for the graphs of order 10. Note the emergence of regular patterns that suggest some kind of regularity in the distribution of phase transitions. But in fact, as shown in S5 Fig, there are triple transitions for a few graphs of order 10. As we said before, these remarks are only apparent because we cannot be sure that there will not exist other transitions for these orders. However, supplementary exact computations for higher values of the fitness *r* (varying from 10 to 2, 000 with step size of 1) corroborate the idea that there is not new transitions in these few graphs. Subject to this caveat, Table 1 gives a detailed account of isothermal and suppressor graphs, as well as graphs exhibiting more than one phase transition. This clarifies the assertion we made above that most graphs of order 10 or less are amplifiers of selection (cf. [9]) or suppressors that become amplifiers from a unique transition phase.

## Discussion

Initially motivated by our interest in the robustness of biological and technological networks against invasion [16], we found in [7] some graph structures suppressing the advantage of mutant individuals occupying their nodes for any fitness value. This seems particularly appealing for biological networks like brain and protein-protein interaction networks, but also in the tumor initiation process within healthy tissues as proposed in [17]. Most graph structures reduce in a very slight amount the advantage of a invading mutant, but some suppression mechanisms could be amplified by repetitive rules (such as those described in [18] and [19] for neuronal networks) involved in the modular architecture of many biological networks.

As explained in Matherials and Methods, we had previously computed the fixation probabilities for all nontrivial connected undirected graphs with 10 or less vertices, totaling 11, 989, 763 graphs, for fitness values varying from 0.25 to 10 with step size of 0.25. Collected data have been published in [14]. Here, we used these data to show that most graphs of order 10 or less are amplifiers of selection. In spite of the complexity of some amplifier structures [4] and the effort required to prove that nature [5,6,8,20,21], this confirms the observation previously made by Hindersin and Traulsen [9] for random graphs of small order. It was also known that some suppressors become amplifiers from some critical fitness value, but here we exhibited new examples (gathered in S1 Fig) whose suppression mechanism is similar to that of clique-wheels studied in [15] and *ℓ*-graphs exhibited in [7]. However, for graphs of order *N* =7, there is another type of transition: we found two amplifiers that become suppressors from critical values *r_c_* ≈ 4.98 and *r_c_* ≈ 6.37. But for larger orders, the phase transitions are even more amazing because some graphs exhibit more than one transition. As before, for *N* =8, these graphs were detected from the barcodes diagram S3 Fig and then identified and represented (see S6 Fig) by using the database [14]. All double transitions are identical of type suppressor/Amplifier/suppressor.

Contrary to the idea that results obtained for weak selection may remain valid out of this case (see [12] and references therein for a discussion about this problem), these observations indicate that the evolutionary regime of a structured population can be completely altered by the graph structure even under constant selection. In our opinion, this has important biological and theoretical implications.

As an analytical computation of the fixation probability for all these graphs does not seem feasible, we adapted the method described in [7] (included in Materials and Methods and whose new code is available from S1 File) to symbolically compute the fixation probabilities of small order graphs with enough symmetries. This allowed us to determine the evolutionary regime of each graph of order 7 or less, providing the exact part of Table 1, which is one of the main results of the paper. The exact value of each transition is included in S1 Table. Thus we found a third amplifier of order 7 with a transition phase at *r_c_* ≈ 24.79, and consequently we were interested in the evolutionary regime of friendship cycles and friendship stars portrayed in S2 Fig when *N* = 7 and *N* = 10.

More generally, for graphs of order *N* = 9, only simple and double transitions are visible in S4 Fig, whereas triple transitions are distinguishable in S5 Fig for order *N* = 10. In the first case, there are 6 graphs with double transitions of type Amplifier/suppressor/Amplifier among a total of 49 graphs with more than one transition. In the second one, we found 19 graphs with triple transitions, all of the same type suppressor/Amplifier/suppressor/Amplifier. All these remarks are part of Table 1. Given the number of graphs of order 9 and 10 having some transition, it would be tedious (and hard without enough symmetries) to determine the exact number of transitions for each graph, so it cannot be excluded the (unlikely) existence of new transitions. However, additional computations for higher values of the fitness *r* (varying from 10 to 2,000 with step size of 1) when the graph has a triple transition seem exclude this possibility.

As we already said, although transitions between different evolutionary regimes were known from [6], [9], [11] and [13], theses results reveal that undirected graphs have a complex and rich evolutionary dynamics, and pose new challenges in computing fixation probabilities and times. Now we know that numerical simulation cannot provide accurate answers to extremal problems on fixation probabilities, nor probably on absorption or fixation times. Our techniques can be easily adapted to determine these times for graphs of small order with enough symmetries. Analyzing the case of disadvantageous mutants (with *r* < 1) is feasible with the same techniques. Also, study the influence of the initial placement of the mutant on the evolutionary outcome for both uniform and temperature initialization (which perhaps is biologically more plausible [8]). In fact, this could clarify better why some suppression mechanisms work effectively. More difficult, however, is to change the updating method, Birth-Death by Death-Birth, converting amplifiers into suppressors and vice versa.

Finally, all barcode diagrams, but specially those of graphs of order 10, show very particular patterns in the distribution of phase transitions. The existence of multiple transitions has been established, but if we think that the fixation probability is a monotonically increasing rational function, this fact is somewhat surprising. What are the features of the graphs leading to multiple phase transitions and their particular distribution is an important issue for future work. Though the real challenge is to translate differential features of suppressors (as has been achieved with some isothermal graphs and amplifiers) into features of their state spaces allowing to analytically compute their fixation probabilities and then understanding their suppression mechanisms.

## Supporting Information

**S1 File. C++ program.** To compute the fixation probabilities of small order graphs for any fitness value *r* > 1, we ran a new C++ program available at **https://bitbucket.org/geodynapp/phasetransition**.

**S1 Table.** Exact phase transitions from order 2 to order 7. Evolutionary state of all connected graphs of order 7 or less whose fixation probability Φ(*r*) = Φ_0_(*r*) for some fitness value *r* > 1.

| ID | Order | Number of cuts | Phase Transition | Type |
| --- | --- | --- | --- | --- |
| 1 | 2 |  |  | Isothermal |
| 1027 | 3 |  |  | Isothermal |
| 526339 | 4 |  |  | Isothermal |
| 527367 | 4 |  |  | Isothermal |
| 135268355 | 5 |  |  | Isothermal |
| 135797775 | 5 |  |  | Isothermal |
| 1580044 | 5 | 1 | 1.78287498125135 | Suppressor/Amplifier |
| 17449355267 | 6 |  |  | Isothermal |
| 17450932235 | 6 |  |  | Isothermal |
| 17585679375 | 6 |  |  | Isothermal |
| 17586207775 | 6 |  |  | Isothermal |
| 17450408974 | 6 |  |  | Isothermal |
| 17586206722 | 6 | 1 | 1.22193837434221 | Suppressor/Amplifier |
| 17584109586 | 6 | 1 | 1.33095684176226 | Suppressor/Amplifier |
| 17586207760 | 6 | 1 | 1.36082690317112 | Suppressor/Amplifier |
| 406338584 | 6 | 1 | 1.87965125079520 | Suppressor/Amplifier |
| 17451461644 | 6 | 1 | 2.82789894107104 | Suppressor/Amplifier |
| 406331401 | 6 | 1 | 4.56890343185820 | Suppressor/Amplifier |
| 1134140852227 | 7 |  |  | Isothermal |
| 1134682454043 | 7 |  |  | Isothermal |
| 1151592840222 | 7 |  |  | Isothermal |
| 1151998655551 | 7 |  |  | Isothermal |
| 1151997603842 | 7 | 1 | 1.01291427227256 | Suppressor/Amplifier |
| 1151863361537 | 7 | 1 | 1.04010585578508 | Suppressor/Amplifier |
| 18127286304 | 7 | 1 | 1.11535825972254 | Suppressor/Amplifier |
| 1151997073410 | 7 | 1 | 1.11712985903186 | Suppressor/Amplifier |
| 1151591804932 | 7 | 1 | 1.13838192561540 | Suppressor/Amplifier |
| 34770286640 | 7 | 1 | 1.15320784414287 | Suppressor/Amplifier |
| 1151456532514 | 7 | 1 | 1.16112279449031 | Suppressor/Amplifier |
| 1151998654496 | 7 | 1 | 1.21650842766890 | Suppressor/Amplifier |
| 1151998101505 | 7 | 1 | 1.23782574145902 | Suppressor/Amplifier |
| 1151998114848 | 7 | 1 | 1.26569832076209 | Suppressor/Amplifier |
| 1151998115845 | 7 | 1 | 1.29194476891885 | Suppressor/Amplifier |
| 1117502071820 | 7 | 1 | 1.32558467744223 | Suppressor/Amplifier |
| 1151862332432 | 7 | 1 | 1.35798530459835 | Suppressor/Amplifier |
| 1151998640129 | 7 | 1 | 1.37227878517787 | Suppressor/Amplifier |
| 1134281915452 | 7 | 1 | 1.47472235317447 | Suppressor/Amplifier |
| 1151998655520 | 7 | 1 | 1.47753803217999 | Suppressor/Amplifier |
| 1151998648321 | 7 | 1 | 1.47874663370948 | Suppressor/Amplifier |
| 1151998654466 | 7 | 1 | 1.49561259905446 | Suppressor/Amplifier |
| 1151998129156 | 7 | 1 | 1.50428354562985 | Suppressor/Amplifier |
| 52487020545 | 7 | 1 | 1.52505989591938 | Suppressor/Amplifier |
| 1151990776850 | 7 | 1 | 1.68966928072836 | Suppressor/Amplifier |
| 1134818766881 | 7 | 1 | 1.77212949793012 | Suppressor/Amplifier |
| 52487027760 | 7 | 1 | 1.80462179670359 | Suppressor/Amplifier |
| 1151998640145 | 7 | 1 | 1.94308343834170 | Suppressor/Amplifier |
| 1151861813254 | 7 | 1 | 1.96627084630835 | Suppressor/Amplifier |
| 1100324302860 | 7 | 1 | 1.98160809559319 | Suppressor/Amplifier |
| 1151994460194 | 7 | 1 | 1.98527430207006 | Suppressor/Amplifier |
| 1134147168268 | 7 | 1 | 2.13899800717826 | Suppressor/Amplifier |
| 1151998648329 | 7 | 1 | 2.16540201237524 | Suppressor/Amplifier |
| 1151863909388 | 7 | 1 | 2.63107033615185 | Suppressor/Amplifier |
| 18127282217 | 7 | 1 | 2.85222067999417 | Suppressor/Amplifier |
| 1151992863778 | 7 | 1 | 2.89510857860393 | Suppressor/Amplifier |
| 1151461244933 | 7 | 1 | 2.95858912425240 | Suppressor/Amplifier |
| 1151058073628 | 7 | 1 | 2.96535184460917 | Suppressor/Amplifier |
| 1151998652455 | 7 | 1 | 3.18958946568448 | Suppressor/Amplifier |
| 1151993934886 | 7 | 1 | 3.38713149404028 | Suppressor/Amplifier |
| 52479149074 | 7 | 1 | 3.41989386804709 | Suppressor/Amplifier |
| 52487012369 | 7 | 1 | 3.51280495555811 | Suppressor/Amplifier |
| 1151996551171 | 7 | 1 | 3.93420590896262 | Suppressor/Amplifier |
| 1151996020755 | 7 | 1 | 4.06745679065459 | Suppressor/Amplifier |
| 1134682447881 | 7 | 1 | 4.77595089988740 | Suppressor/Amplifier |
| 1151592835082 | 7 | 1 | 4.98811715947310 | Amplifier/Suppressor |
| 1151864430605 | 7 | 1 | 5.08434009504437 | Suppressor/Amplifier |
| 1151998648333 | 7 | 1 | 5.17611822443887 | Suppressor/Amplifier |
| 1151860745228 | 7 | 1 | 6.37125003382654 | Amplifier/Suppressor |
| 1151994973210 | 7 | 1 | 6.86047703514633 | Suppressor/Amplifier |
| 1134549822492 | 7 | 1 | 6.88630847258735 | Suppressor/Amplifier |
| 1134416132124 | 7 | 1 | 8.93833010810699 | Suppressor/Amplifier |
| 1151864426517 | 7 | 1 | 10.45645176381780 | Suppressor/Amplifier |
| 1151997074462 | 7 | 1 | 13.61188814438320 | Suppressor/Amplifier |
| 1151592837126 | 7 | 1 | 24.79702019687890 | Amplifier/Suppressor |
| 1151998652423 | 7 | 1 | 25.47101313451720 | Suppressor/Amplifier |
| 1151590742058 | 7 | 1 | 77.28669194491090 | Suppressor/Amplifier |
| 1151997057027 | 7 | 1 | 79.15367807274810 | Suppressor/Amplifier |
| 1151592839194 | 7 | 1 | 85.75032790283100 | Suppressor/Amplifier |

**S2 Table.**
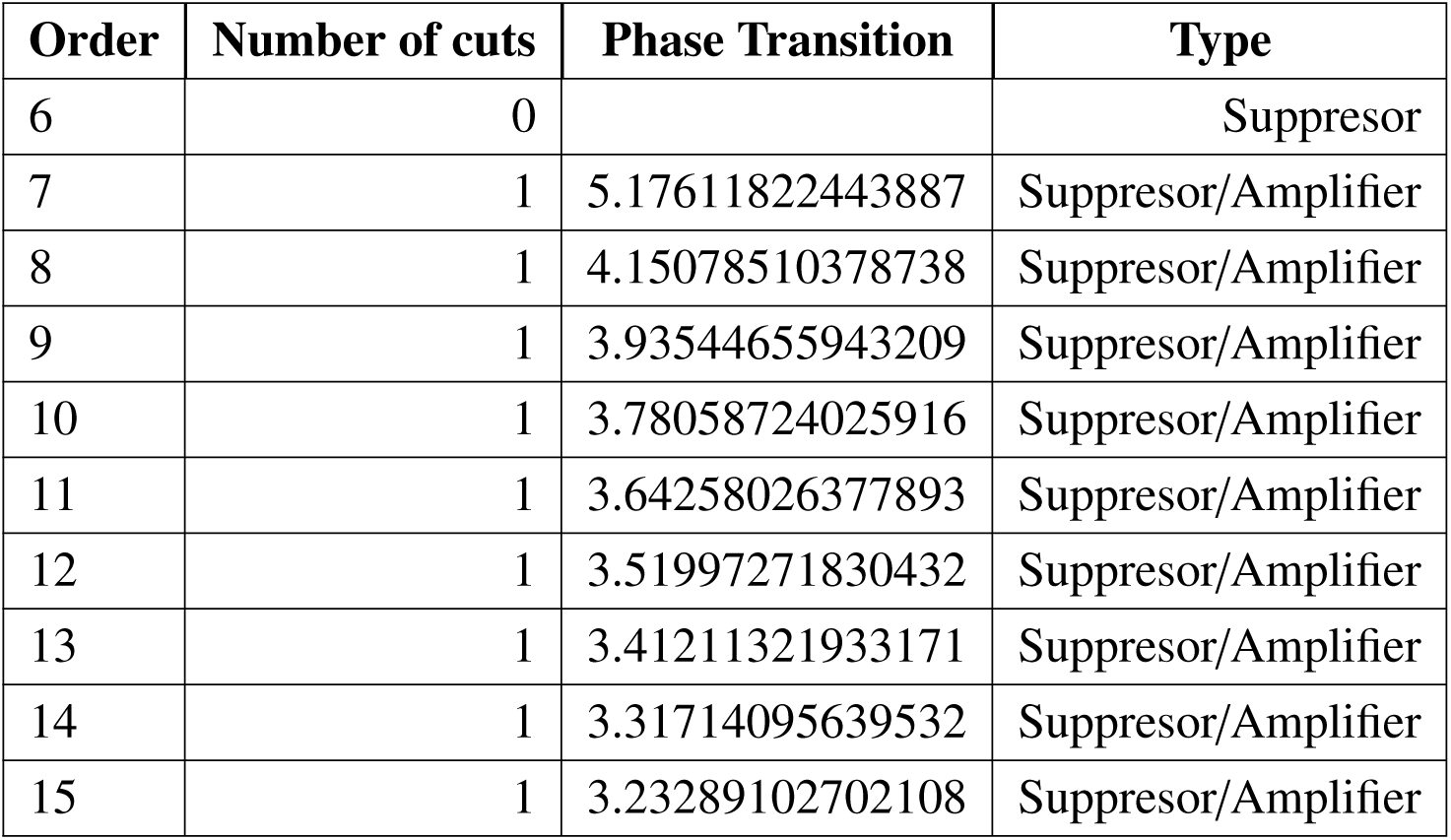
Exact phase transitions for graphs
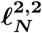
from order 6 to order 15.

**S1 Fig.**
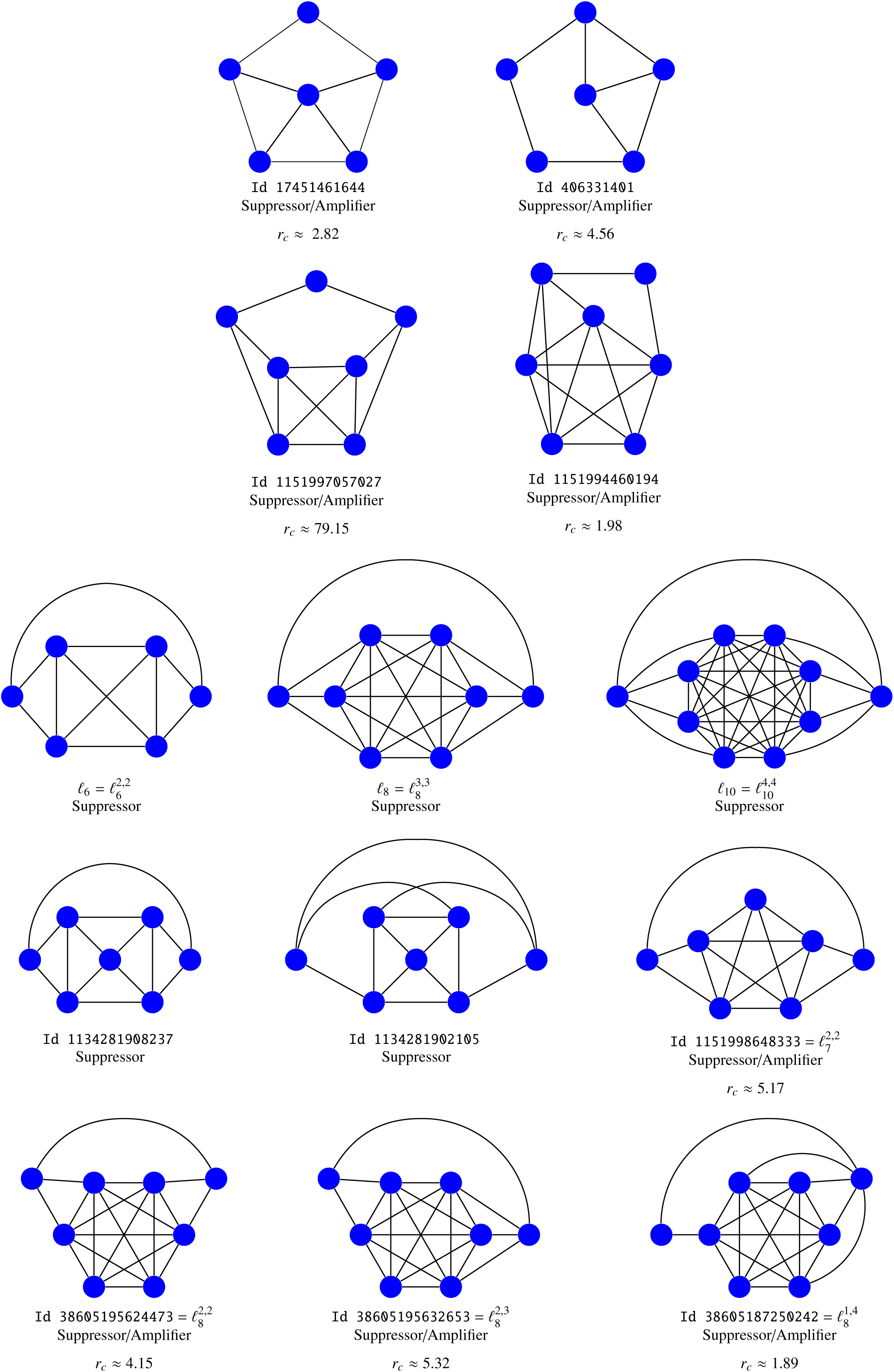
Suppressors for weak selection (and beyond). All the figures representing suppressors are gathered with indication of their identification numbers, names, regimes and transitions.

**S2 Fig.**
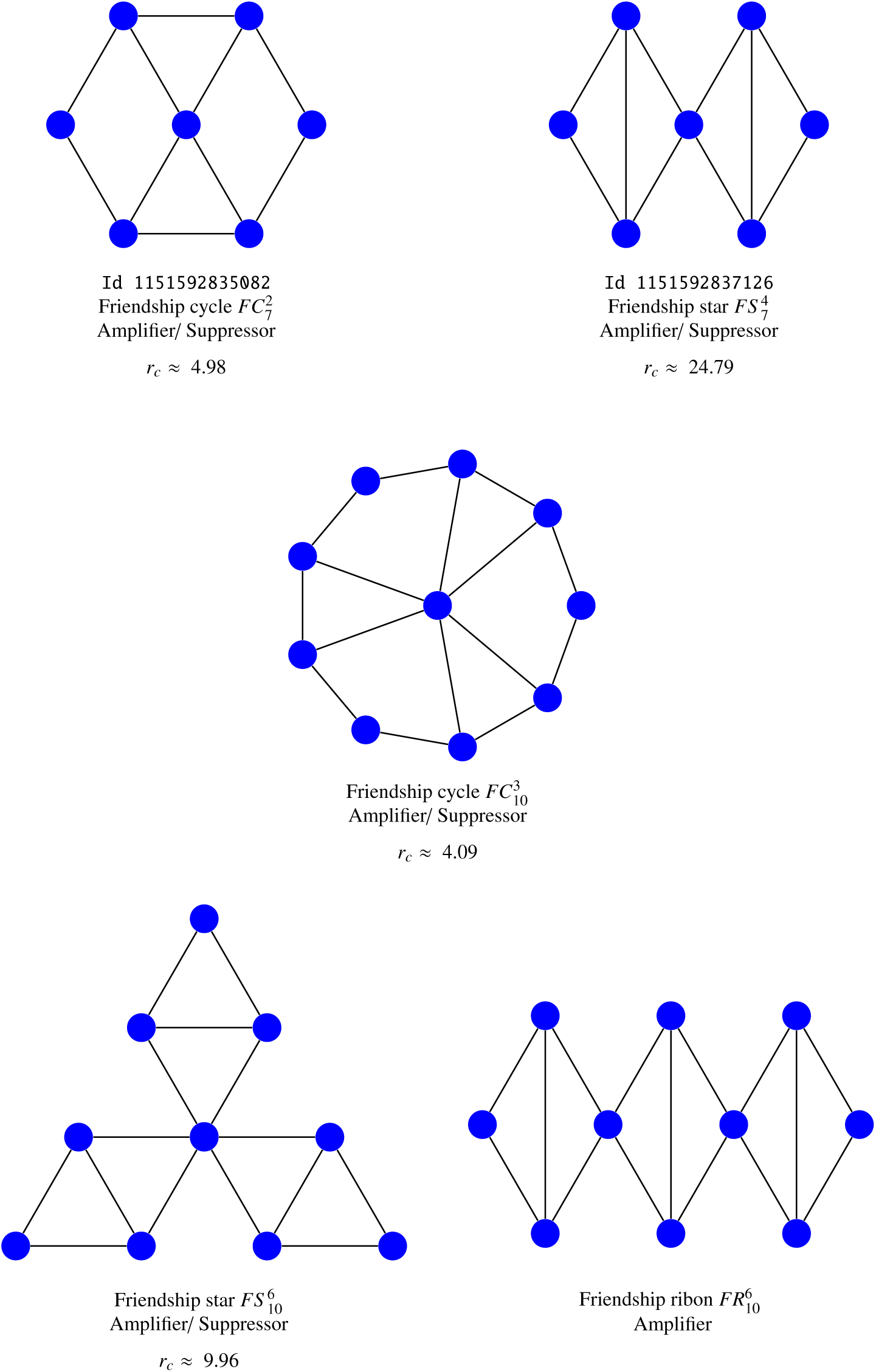
Friendship cycles, ribbons and stars. Some friendship cycles
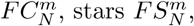
and ribbons
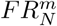
of order *N* = 7 and 10 are compared.

**S3 Fig.**
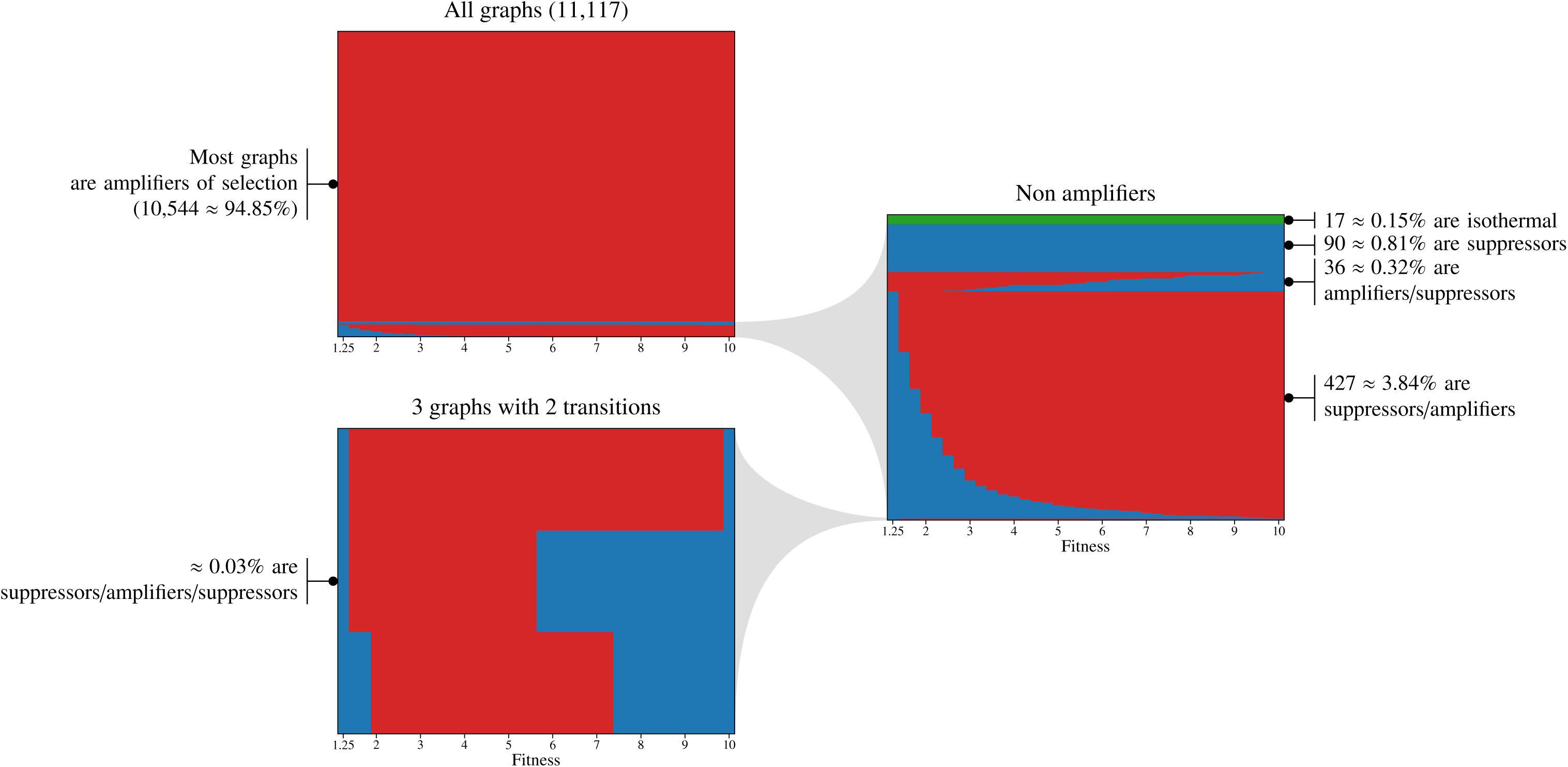
Barcodes describing transition phases of graphs of order 8. Each horizontal line corresponds to a graph, and color represents the evolutionary regime for the given fitness: blue color corresponds to suppressor regime and red color to amplifier regime.

**S4 Fig.**
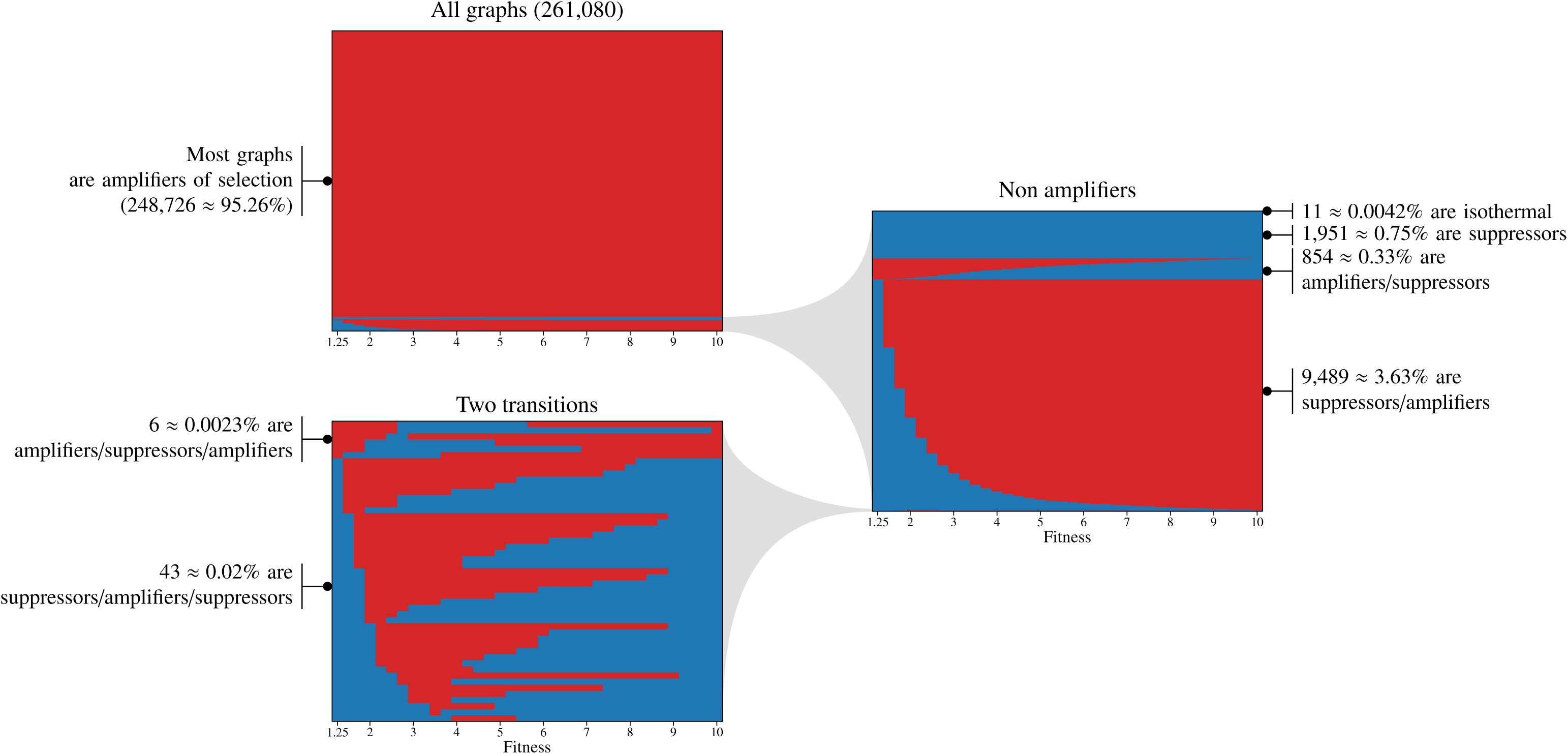
Barcodes describing transition phases of graphs of order 9. Each horizontal line corresponds to a graph, and color represents the evolutionary regime for the given fitness: blue color corresponds to suppressor regime and red color to amplifier regime.

**S5 Fig.**
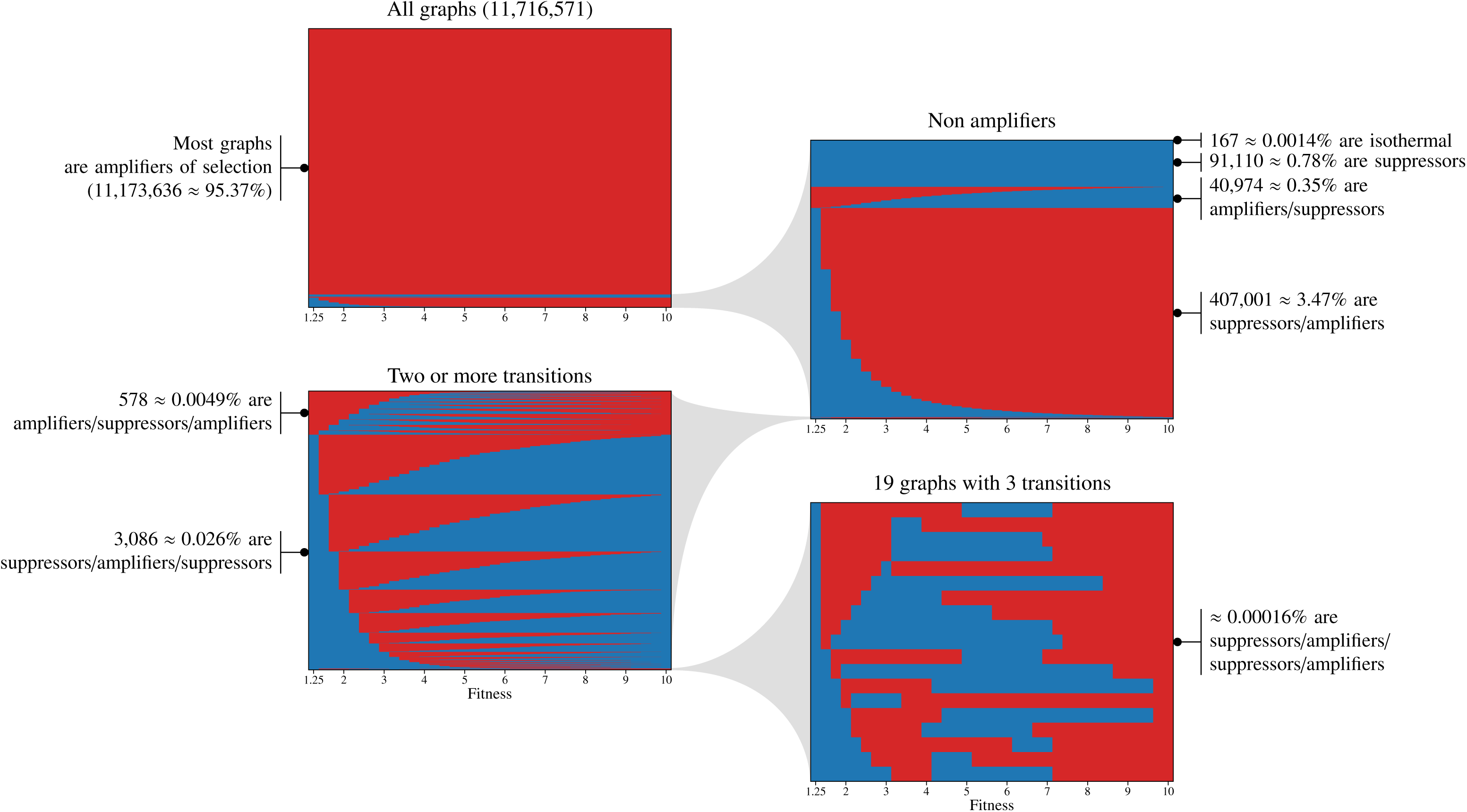
Barcodes describing transition phases of graphs of order 10. Each horizontal line corresponds to a graph, and color represents the evolutionary regime for the given fitness: blue color corresponds to suppressor regime and red color to amplifier regime.

**S6 Fig.**
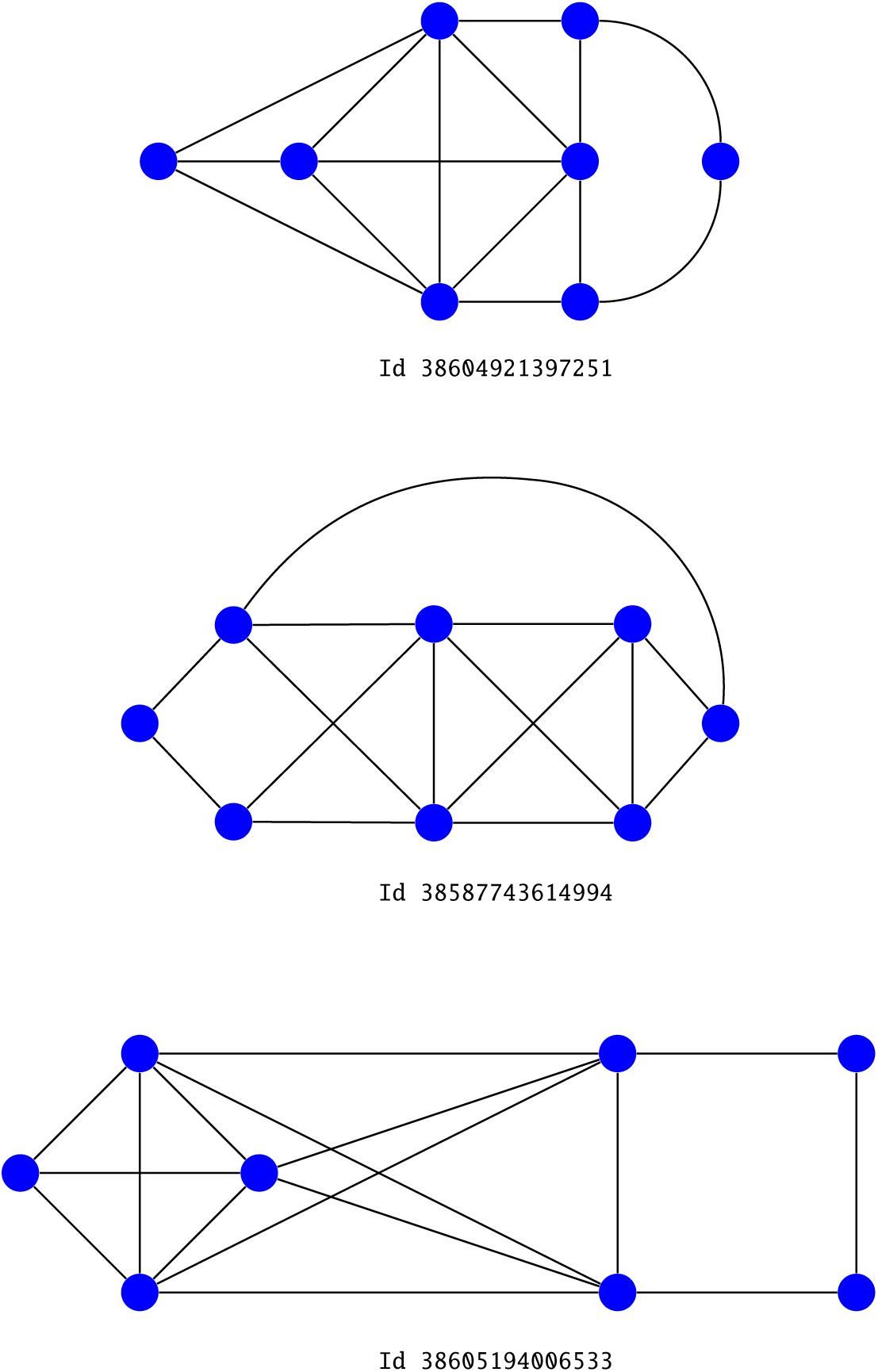
Graphs of order 8 with a double transition of type Suppressor/Amplifier/Suppressor.

## References

1. Moran PAP. Random processes in genetics. Proc Cambridge Philos Soc. 1958;54:60–71.

2. Maruyama T. On the fixation probability of mutant genes in a subdivided population. Genetical Research. 1970;15(2):221–225. doi:10.1017/S0016672300001543.

3. Maruyama T. A simple proof that certain quantities are independent of the geographical structure of population. Theoretical Population Biology. 1974;5(2):148–154. doi:10.1016/0040-5809(74)90037-9.

4. Lieberman E, Hauert C, Nowak MA. Evolutionary dynamics on graphs. Nature. 2005;433(7023):312–316.

5. Monk T, Green P, Paulin M. Martingales and fixation probabilities of evolutionary graphs. Proceedings of the Royal Society of London A: Mathematical, Physical and Engineering Sciences. 2014;470(2165). doi:10.1098/rspa.2013.0730.

6. Broom M, Rychtář J, Stadler BT. Evolutionary dynamics on graphs—the effect of graph structure and initial placement on mutant spread. J Stat Theory Pract. 2011;5(3):369–381. doi:10.1080/15598608.2011.10412035.

7. Alcalde Cuesta F, González Sequeiros P, Lozano Rojo A. Suppressors of selection. PLOS ONE. 2017;12(7):1–11. doi:10.1371/journal.pone.0180549.

8. Adlam B, Chatterjee K, Nowak MA. Amplifiers of selection. Proceedings of the Royal Society of London A: Mathematical, Physical and Engineering Sciences. 2015;471(2181). doi:10.1098/rspa.2015.0114.

9. Hindersin L, Traulsen A. Most Undirected Random Graphs Are Amplifiers of Selection for Birth-Death Dynamics, but Suppressors of Selection for Death-Birth Dynamics. PLoS Comput Biol. 2015;11(11):1–14. doi:10.1371/journal.pcbi.1004437.

10. Kaveh K, Komarova NL, Kohandel M. The duality of spatial death–birth and birth–death processes and limitations of the isothermal theorem. Royal Society Open Science. 2015;2(4). doi:10.1098/rsos.140465.

11. Alcalde Cuesta F, González Sequeiros P, Lozano Rojo Á, Vigara Benito R. An Accurate Database of the Fixation Probabilities for All Undirected Graphs of Order 10 or Less. In: Rojas I, Ortuño F, editors. Bioinformatics and Biomedical Engineering: 5th International Work-Conference, IWBBIO 2017, Granada, Spain, April 26–28, 2017, Proceedings, Part II. Cham: Springer International Publishing; 2017. p. 209–220.

12. Wu B, García J, Hauert C, Traulsen A. Extrapolating Weak Selection in Evolutionary Games. PLOS Computational Biology. 2013;9(12):1–7. doi:10.1371/journal.pcbi.1003381.

13. Choi JO, Yu U. Fixation probability on clique-based graphs. Physica A: Statistical Mechanics and its Applications. 2018;492:2129–2135. doi:https://doi.org/10.1016/j.physa.2017.11.131.

14. Alcalde Cuesta F, González Sequeiros P, Lozano Rojo Á, Vigara Benito R. An accurate database of the fixation probabilities for all undirected graphs of order 10 or less, Mendeley Data, v2; 2017. Available from: https://data.mendeley.com/datasets/587bnf6mt3/2.

15. Mertzios GB, Nikoletseas S, Raptopoulos C, Spirakis PG. Natural models for evolution on networks. Theoretical Computer Science. 2013;477(Supplement C):76–95. doi:10.1016/j.tcs.2012.11.032.

16. Alcalde Cuesta F, González Sequeiros P, Lozano Rojo Á. Exploring the topological sources of robustness against invasion in biological and technological networks. Scientific Reports. 2016;6:20666 EP –. doi:10.1038/srep20666.

17. Hindersin L, Werner B, Dingli D, Traulsen A. Should tissue structure suppress or amplify selection to minimize cancer risk? Biology Direct. 2016;11(1):41. doi:10.1186/s13062-016-0140-7.

18. Perin R, Berger TK, Markram H. A synaptic organizing principle for cortical neuronal groups. Proceedings of the National Academy of Sciences. 2011;108(13):5419–5424. doi:10.1073/pnas.1016051108.

19. Azulay A, Itskovits E, Zaslaver A. The C. elegans Connectome Consists of Homogenous Circuits with Defined Functional Roles. PLOS Computational Biology. 2016;12(9):1–16. doi:10.1371/journal.pcbi.1005021.

20. Voorhees B, Murray A. Fixation probabilities for simple digraphs. Proceedings of the Royal Society of London A: Mathematical, Physical and Engineering Sciences. 2013;469(2154). doi:10.1098/rspa.2012.0676.

21. Shakarian P, Roos P, Moores G. A novel analytical method for evolutionary graph theory problems. Biosystems. 2013;111(2):136–144. doi:10.1016/j.biosystems.2013.01.006.

